# Improving the predictive performance of CLUE-S by extending demand to land transitions: the trans-CLUE-S model

**DOI:** 10.1101/2023.01.10.523486

**Authors:** Diogenis A. Kiziridis, Anna Mastrogianni, Magdalini Pleniou, Spyros Tsiftsis, Fotios Xystrakis, Ioannis Tsiripidis

**Author notes:** Corresponding author: Diogenis A. KIZIRIDIS, Department of Botany, School of Biology, Aristotle University of Thessaloniki, Thessaloniki, Greece, PC 54124.

## Abstract

The CLUE-S model is a popular choice for modelling land use and land cover change from local to regional scales, but it spatially allocates the demand for only the total cover of each land class in the predicted map. In the present work, we introduce a CLUE-S variant that allocates demand at the more detailed level of land type transitions, the trans-CLUE-S model. We implemented this extension algorithmically in R, without the need of new parameters. By processing each row of the land transition matrix separately, the model allocates the demand of each land category’s transitions via the CLUE-S allocation routine for only the cells which were of that category in the map of the previous time step. We found that the trans-CLUE-S model had half the total and configuration disagreement of the CLUE-S predictions in an empirical landscape, and in simulated landscapes of different characteristics. Moreover, the trans-CLUE-S performance was less sensitive to the number of environmental predictors of land type suitability for allocating demand. Although trans-CLUE-S is computationally more demanding due to running a CLUE-S allocation for each land class, we appended the solution of a land-use assignment optimisation problem that facilitates the convergence and acceleration of allocation. We additionally provide R functions for: CLUE-S variants at other levels of demand resolution; random instead of environment-based allocation; and for simulating landscapes of desired characteristics. Our R code for the models and functions can contribute to more reproducible, transparent and accurate modelling, analysis and interpretation of land cover change.

**Highlights:**

- The trans-CLUE-S model employs demand at the finer level of land type transitions
- The trans-CLUE-S predictions were twice more accurate than the CLUE-S model’s
- The trans-CLUE-S accuracy was less dependent on the amount of environmental data
- Algorithmic addition of a land assignment task enabled and sped up full convergence
- R code is provided for our models and auxiliary functions

## 1. Introduction

Land use and cover (LUC) change is a major anthropogenic driver of environmental change from local to global scales (Verburg et al., 2015). Since field experiments are commonly prohibitive, models of LUC change enable the mathematical and *in silico* experimentation with underlying socioeconomic, biophysical and other environmental conditions at different spatial scales (van Vliet et al., 2016). According to Mas et al. (2014), such LUC models are characterised on the basis of their temporal dimension (dynamic versus static), spatial dimension (spatial versus non-spatial), inference about pattern–process relations (inductive versus deductive), and rules of LUC change (pattern-based versus agent-based). The development of new models, and the extension or hybridisation of existing ones, hence constitute an active area of research which improves our knowledge, forecasts and guidance to decision-making related to LUC change (Ren et al., 2019). With the aim to improve predictions at a relatively small extra cost in model complexity and computational resources, the present work developed novel variants of the dynamic, spatial, inductive and pattern-based model called CLUE-S (Verburg et al., 2002).

The Conversion of Land Use and its Effects at Small regional extent (CLUE-S) is a popular choice for modelling LUC change in finer spatial resolution (map grid cells of ≲ 1 km side length), and from local to regional scales (Ren et al., 2019). The CLUE-S, as other spatial models, is constituted by two main parts: the non-spatial part of demand in land type cover, and the spatial part of demand allocation. The basic task of the model, then, is the spatial allocation of this pre-specified demand. Regarding the demand part, CLUE-S requires the total number of cells which will be covered by each land type in the future map to be predicted (Ren et al., 2019; Verburg et al., 2002).

These land type sums can be taken by a transition matrix built from a past time interval, by extrapolation of the historical time series of land type sums to the future, or by economic models (Fuchs et al., 2013; Moulds et al., 2015). Regarding the allocation part, CLUE-S requires statistical models which relate the probability of occurrence of each land type to environmental conditions (Verburg et al., 2002). These statistical models are fitted on historical data, and are employed for selecting where to spatially allocate the land type demand in the map of the future time point for which environmental conditions will be given. Being of inductive type, CLUE-S allocation is hence based on the so-called “suitability” of the cells to each land type. The original CLUE-S software employs a separate logistic regression model of environmental suitability for each land type (Verburg et al., 2002).

With these statistical models of suitability, the future demand of each land type is consequently allocated in a top-down way by the CLUE-S model (Ren et al., 2019). The CLUE-S allocation setting has hence its strengths and weaknesses. On the one hand, it requires more easily acquired environmental data, for only the building of the statistical models of suitability, since mechanistically involved socioeconomic forces, such as irrigation and wood demand from a broader scale which can contribute to LUC change at the focal scale (Harrison et al., 2016; Holman et al., 2017), are implicitly assumed and incorporated to the estimation of demand. Additionally, the whole LUC system’s functioning can be explicitly taken into account by the CLUE-S, simulating the competition of multiple LUC types for allocation, with the use of reproducible statistical models of suitability (Ren et al., 2019; Rosa et al., 2014). On the other hand, these two strengths of CLUE-S have their respective drawbacks. First, the implicit accounting of socioeconomic factors, mechanisms and actors in the projected demand, and the statistical models of suitability allocation of demand, can limit the interpretability of the CLUE-S predictions. Second, a greater amount of historical data on LUC trajectories and environmental conditions might be required, due to this heavier reliance on statistical models of demand and suitability. All these strengths and weaknesses of CLUE-S make it particularly suitable for simulating LUC change, and for analysing a wide range of study areas and scenarios, especially for regions which lack socioeconomic data of broader scale and finer quality (Ren et al., 2019).

Besides the aforementioned weaknesses, the demand of the current CLUE-S model is limited to the level of land type sums in the future map for prediction. This is less detailed than in other spatially explicit models which have their demand at the finer level of land type transitions. In specific, such models expect as demand the number of cells for each land type transition from the latest map to the new map to be predicted (such models are reviewed in: Mas et al., 2014; Ren et al., 2019). CLUE-S has two features which might compensate for the lack of demand information at the level of land type transitions: (1) five transition rules which can prohibit transitions in space and time; and (2) land type elasticity which can specify how elastic each land type is to change. However, the adoption of strict enough transition rules can prevent the allocation algorithm from meeting the specified demand (Moulds et al., 2015). Moreover, these two features, which contribute considerably to model complexity with the adoption of extra parameters to tune, are not always straightforward to parameterise, commonly requiring expert judgement (Mas et al., 2014). The latter authors have hence noted that the CLUE-S demand at the resolution of land type sums is a feature which can be easily replaced by more informative approaches, e.g. demand at the resolution of land type transitions.

In the present work, we introduce four novel CLUE-S variants of increasing demand resolution, the most detailed of which is at the resolution of land type transitions. For simplicity, we focused on the comparison between the original CLUE-S and our most detailed variant with demand for land type transitions. We show that this detailed variant that we call trans-CLUE-S is twice more accurate than the original CLUE-S in the prediction of an empirical landscape, and of numerous simulated landscapes with a wide range of landscape characteristics. Additionally, trans-CLUE-S is less sensitive than CLUE-S to the number of environmental suitability predictors employed for the allocation of demand. This is important because environmental information, especially for the future, is commonly limited or of high variability, and trans-CLUE-S can hence provide more robust LUC projections. The R functions for running the models, together with other functions—such as a function for building simulated landscapes of desired characteristics, and a function for facilitating the convergence and acceleration of any CLUE-S-based allocation— are freely available (https://doi.org/10.6084/m9.figshare.21014869.v1).

## 2. Materials and methods

CLUE-S, and therefore our introduced variants as well, require mainly two types of input data: (1) categorical maps of observed LUC of the study area at historical time points, i.e. each cell covered by a single land type at each time point; and (2) biophysical, socioeconomic and other environmental conditions of the study area for the corresponding cells and time points. As with other empirical models, a spatial model’s preparation procedure has three phases (Pontius et al., 2004): parameterisation (tuning the model); prediction (running the model); and validation (assessing the model’s predictions). A model is usually parameterised by input data from all time points except the last one. The model is then run to predict the map of the last time point. Finally, the predicted map is validated against the observed reference map of the last time point (Pontius et al., 2004). After enough parameterisation–prediction–validation rounds leading to a satisfactory model, the model is parameterised by using data also from the last map at the time point *t* = 1 (map 1 hereafter), and it is called to predict the LUC map 2 at a future time step *t* = 2, given the environmental conditions at *t* = 2.

CLUE-S and other spatial models are parameterised mainly at their two core parts: the non-spatial component of land type demand, and the spatial component of suitability-based demand allocation. For the demand part, a frequent starting point is the estimation of a transition matrix built via the cross-tabulation of land type frequencies between two consecutive maps of a historical time interval before the desired prediction interval of map 1–map 2, e.g. between the map 0 at an adjacent, previous time step *t* = 0, and the latest historical map 1. The entries of the transition matrix denote the number of cells which transitioned from the land types in the rows (map 0) to the land types in the columns (map 1). The row and column sums hold the total number of cells covered by each land type in map 0 and map 1, respectively. By dividing a matrix entry by its row’s total number of cells, we get a Markov matrix of transition probabilities, summing up to one in each matrix row. For a land type in its row, the Markov matrix can then be used to project the demanded transitions in map 2 from map 1, by multiplying the row’s entries by the total number of cells covered by the land type in map 1. If the map 1–map 2 time interval is not of the same duration as the map 0–map 1 interval, there exist techniques for projecting the transition matrix of the map 0–map 1 interval to the duration of the map 1–map 2 interval (Eastman and He, 2020; Takada et al., 2010).

For the suitability part, the original CLUE-S software employs one logistic regression model for the presence of each land type, but binomial classification trees and binomial classification random forest models for suitability have been added in an R implementation of CLUE-S (Moulds et al., 2015). We herein extended this R implementation by adding the possibility of multinomial classification models, which provide a neater and computationally cheaper alternative (one statistical model for all land types), also shown to perform better in the allocation of demand (Lin et al., 2014). Additionally, we provide random (null, uninformed) suitability allocation versions for the CLUE-S model and our new demand variants. The suitability of each cell to each land type can be predicted by statistical models based on environmental predictors for time step *t* = 2, or it can be random. In the case of environmental suitability, the model variants were deterministic, i.e. returned the same prediction if run multiple times. The only source of stochasticity could be the random selection of a land type in case a cell had equal suitability for two or more land types. We ruled out this source of stochasticity by always selecting the last land type in the list of ties in suitability.

In the present work, although we did not parameterise and hence did not use any extra setting, the R implementations of CLUE-S and our variants preserve the extra features of transition rules and elasticity from the R implementation we were based upon, and from the original CLUE-S software (Moulds et al., 2015; Verburg et al., 2002). Regarding the five transition rules, it would be possible to prevent a cell’s land type from changing: (1) completely throughout space and time, e.g. in the case of urban land which is unlikely to turn to farmland due to the high cost of initially building houses and streets; (2) if it has not persisted for a minimum number of time steps; (3) if it has changed for a maximum number of steps; (4) outside the land type’s defined spatial neighbourhood; and (5) in specific localities throughout time.

### 2.1. Allocating demand with CLUE-S and three new variants

We first present the core algorithm of the original CLUE-S allocation, and then of our variants at the highest level of land type transitions, at the level of land type sums and persistence, and finally the one of no demand information:

#### 1. CLUE-S

The original CLUE-S model requires the demand DMD_*lt*_ at the level of sum for each land type *lt* in map 2 (Fig. 1). The sums can be taken from the column sums of the map 1–map 2 transition matrix. Additionally, CLUE-S requires for each cell *c* its suitability SUIT_*c,lt*_ for each land type. At the first iteration loop, the allocation algorithm assigns to each cell the land type LT_*c*_ with the highest suitability. This initial allocation can disagree with the expected demand if the frequency of cells FREQ_*lt*_ assigned to a land type deviates from the land type’s demand DMD_*lt*_ for more than a pre-specified deviation limit (in number of cells). If the allocated cells are more than demanded, FREQ_*lt*_ > DMD_*lt*_, SUIT_*c,lt*_ to this land type is decreased for all cells; if the allocated cells are fewer than demanded, FREQ_*lt*_ < DMD_*lt*_, SUIT_*c,lt*_ to this land type is increased for all cells. In both cases, the alteration of SUIT_*c,lt*_ is proportional to the distance between FREQ_*lt*_ and DMD_*lt*_, i.e. scaled by a factor. If this scaling factor is large enough, we observed that the new distance between FREQ_*lt*_ and DMD_*lt*_ can become greater than the distance of the previous iteration, forbidding any allocation convergence. We hence enabled the self-adjustment of the factor, by halving its value any time the new distance between FREQ_*lt*_ and DMD_*lt*_ was greater than the previous iteration’s. This had the additional benefit of speeding up convergence, since the alteration of SUIT_*c,lt*_ with a larger factor is more drastic at the initial iterations of larger deviation from demand, but alteration becomes finer as the deviation limit is approached. The iterative alteration of suitability stops when demand is satisfied to the desired deviation limit for all land types, or if a certain number of iterations is reached (Fig. 1).

**Fig. 1.**
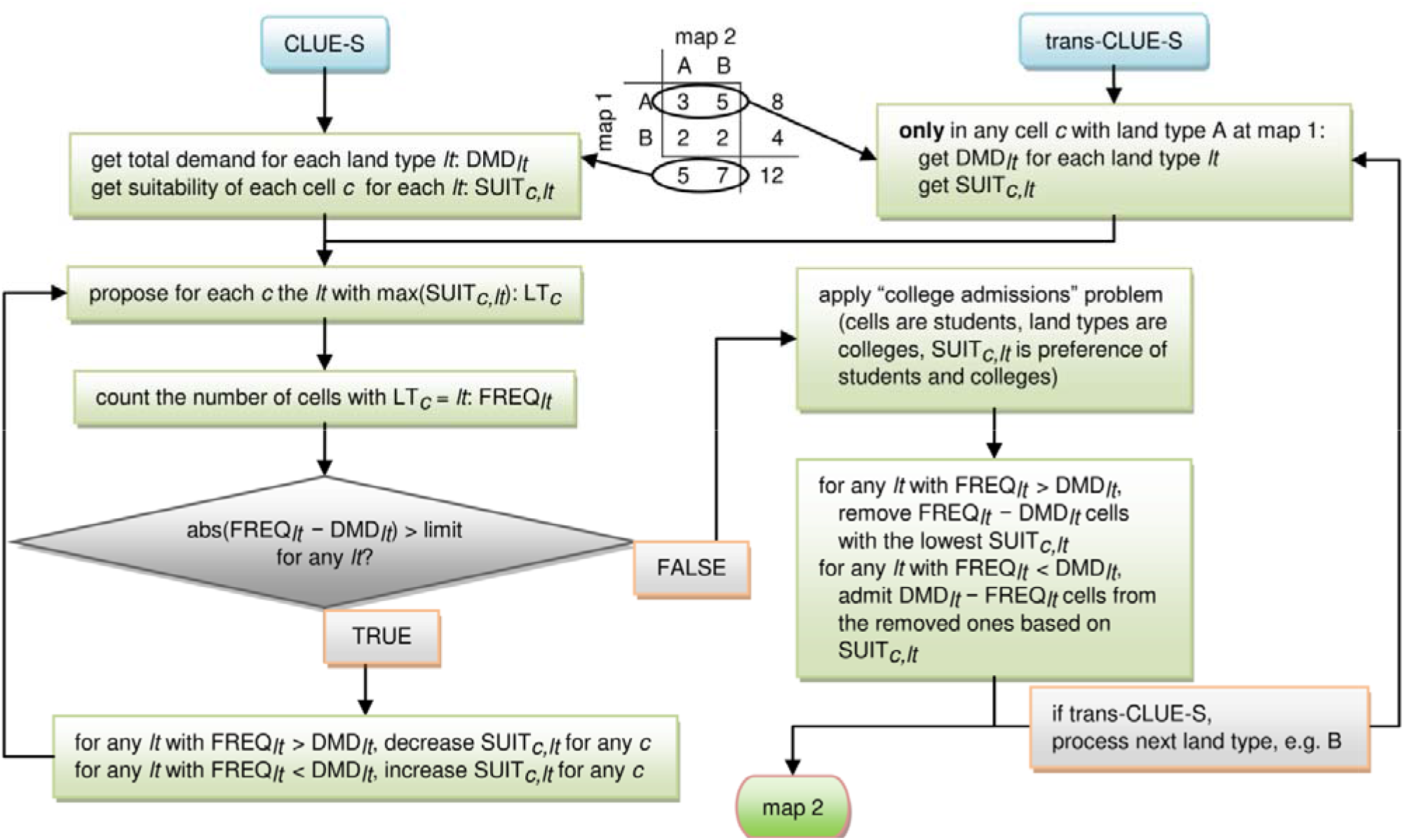
Flowcharts for the core algorithms of the original CLUE-S model (left branch), and of its extended “trans-CLUE-S” variant introduced in this work (right branch). The flowcharts for the two models focus on one time step, i.e. predicting a map at time step *t* = 2 (map 2) given a map at *t* = 1 (map 1). An example transition matrix with two land types is given at the top. Note that the trans-CLUE-S model essentially runs a separate CLUE-S allocation loop for each land type, within each row of the transition matrix. We have also appended our college admissions function, which facilitates the convergence and acceleration of allocation after a deviation limit between allocated and demanded number of cells has been reached by the end of the CLUE-S allocation routine.

#### 2. trans-CLUE-S

This variant requires demand at the level of land type transitions. It essentially runs a CLUE-S allocation within each land type in map 1 separately, instead of the entire landscape. That is, the simulation concentrates on the cells which were of a focal land type in map 1 (the total number of the focal cells is the corresponding row’s sum in the transition matrix). The demand for a focal land type in map 2 is taken from the land type’s row in the transition matrix (Fig. 1). We then know how many cells will persist (the row’s entry at the main diagonal), and how many cells will turn to other land types (off-diagonal entries). As in CLUE-S, we altered iteratively the suitability of the deviated land types until demand is satisfied to a desired deviation for the focal land type (Fig. 1). The same routine is executed for the cells of the other land types in map 1.

#### 3. Land type sums and persistence

This variant is a hybrid between CLUE-S and trans-CLUE-S. Initially, it focuses on the transition matrix’s main diagonal which holds the number of persisting cells for each land type. From the cells in map 1 of each land type, the algorithm selects the cells with the highest suitability for that land type, and flags them as persistent. Next, it runs a CLUE-S allocation to the rest of the cells, but after subtracting the main diagonal entries from the demand of the column sums.

#### 4. No demand

This variant merely assigns to each cell the land type with the highest suitability. It essentially returns a plain suitability map for time step *t* = 2.

To facilitate the convergence of the allocation algorithm after a desired deviation from demand was reached, we appended the solution of a college admissions problem to the execution of the three first variants above (Fig. 1). The college admissions problem formulates the process of students applying to preferred colleges, whereas colleges admit the most preferred students (Gale and Shapley, 1962). Correspondingly, in our LUC setting, cells are the students, land types are the colleges, and the suitability is the preference of the student (cell) for each college (land type). At the same time, we assumed that a land type will cover a cell with high enough suitability, like assuming that a college simply prefers more a student who is more willing to study in that college. Starting with the allocated land cover which deviated from demand by the end of the CLUE-S iterations, our procedure could satisfy demand completely by removing from any land type with excessive cells the ones with the lower suitability to that land type, and by relocating them to any cell-deficient land type they were more suitable for (Fig. 1). This LUC-equivalent implementation of the college admissions problem was formulated and executed by the relevant func ion of the R package “matchingR” (Tilly and Janetos, 2021).

### 2.2. Applying the models to an empirical reference landscape

The empirical reference landscape was a circular study area of 6 km diameter (28.3 km^2^ cover), located at the mountainous region of Pindus in northwestern Greece. The region’s dominant woody taxa classify it to the “thermophilous deciduous oaks” vegetation formation (Bohn et al., 2007). We have mapped the area’s LUC for years 1945, 1970, 1996 and 2015 (Kiziridis et al., 2022). The mapping focused on five land types which are broad steps of progressive vegetation succession: farmland, grassland, open-scrub, closed-scrub and forest. Settlements and bodies of water were excluded, leading to a mapped 27.4 km^2^ cover which is smaller than the circle’s cover. Visual interpretation of orthoimages was used to identify LUC, before proceeding to vectorisation, and rasterisation at 25 m resolution. We used the LUC maps of 1996 and 2015 as the respective reference maps 1 and 2. The largest LUC changes in this 1996–2015 time interval were the decrease of farmland from 16.3 to 6.1% of the mapped cover, the increase of grassland from 7.1 to 12.3%, and the increase of forest from 62.4 to 70.2%. Additionally, the majority of land type transitions occurred towards the progressive successional pathway.

Regarding the environmental conditions, the region’s climate is temperate, belonging to the “Csa” hot-summer Mediterranean type according to the Köppen–Geiger classification scheme (Peel et al., 2007). The study area has an altitude ranging from 390 to 1203 m, and a slope ranging from 0–48°. The wider region had a history of low-intensity agriculture and transhumant livestock grazing until the 1940s, but abandonment of farmlands and grasslands thenceforth has commonly led to vegetation succession and afforestation (Kiziridis et al., 2022; Zomeni et al., 2008). During 1996–2015, at the study area in particular, the population density median decreased from 17.3 to 12.1 inhabitants km^−2^, and the livestock density median decreased from 135.9 to 51.9 small grazing livestock units km^−2^. More details about the empirical reference landscape can be found in Kiziridis et al. (2022).

For building the statistical model of environmental suitability, we used raster data of 25 m resolution from 13 biophysical and socioeconomic predictors from all study years except the last year 2015. Six of the environmental predictors of land type occurrence were fixed in time (altitude, slope, northness, eastness, presence of silicate parent rock and presence of flysch parent rock), whereas seven could vary in time (temperature annual range, temperature seasonality, annual precipitation, precipitation seasonality, population density, livestock density and distance to the nearest settlement). We related land type occurrence to the environmental predictors via random forest, multiclass classification modelling using a balanced subset of the data (Biau and Scornet, 2016). We employed the “ranger” method to train the model with the R package “caret” (Kuhn et al., 2021), by using 10-fold cross-validation, and fine-tuning two model hyperparameters. All different combinations were tried for these two hyperparameters of the split rule (“gini” or “extratrees”) and of the number of randomly selected predictors to employ at each tree split (two, half or all of the predictors). We kept the paired combination of hyperparameter values that lead to the maximum performance in classification according to the “Accuracy” and “Kappa” metrics. The hyperparameter of minimum number of observations in any terminal node of any individual tree was fixed to one. Finally, the hyperparameter of the tree number was fixed to 1000 trees, since random forest performance is robust to their count (Probst and Boulesteix, 2018).

We applied the CLUE-S and the trans-CLUE-S models to the empirical landscape, to predict the map of year 2015 (map 2) under either environmental or random suitability allocation. For reference, we additionally ran the no-demand version, i.e. simply predicting the environmental suitability with the random forest model, or the random suitability in map 2. The three demand versions and two suitability modes of allocation resulted to 3 × 2 = 6 model versions to compare in total. Where applicable, the demand for the models was taken by the reference transition matrix of the 1996–2015 time interval (reference maps 1–2). For comparability and simplicity, no model version adopted any constraints in the five transition rules or in the elasticity settings.

### 2.3. Applying the models to simulated reference landscapes

We aimed to compare the models in a wide variety and large number of different landscapes, and we hence decided to create such landscapes *in silico*. The advantage of using simulated reference landscapes, instead of a collection of empirical ones, is that we could systematically vary specific characteristics of the simulated landscapes, and for a large enough number of replicates to strengthen our inference. Thus, to infer possible weaknesses and strengths of the compared models in landscapes of different characteristics, we varied the three following landscape parameters:

#### 1. The number of land types

This parameter was the number of land types in reference maps 1 and 2. For simplicity, we restricted all land types to occur in both reference maps 1 and 2, covering each map with at least five pixels each. Given the commonly mapped numbers of land types in different applications of empirical models (Pontius et al., 2008; van Vliet et al., 2016), we created landscapes with five different numbers of land types: {3, 6, 9, 12, 15}.

#### 2. The spatial aggregation

This parameter controlled the spatial clustering of the land types in reference map 1. It was taken as equivalent to the relative mutual information, a landscape metric quantifying the information that a focal raster cell’s land type provides for the prediction of an adjacent cell’s land type, normalised by the overall land type diversity (Nowosad and Stepinski, 2019). Thus, relative mutual information functions as an unbiased measure of spatial clustering, comparable among landscapes with different numbers of land types, and with different evenness of land type cover. Relative mutual information approaches its minimum value of zero when the land type prediction of a neighbouring cell cannot be better than random, and attains its maximum value of one when the landscape is of one land type (herein rescaled to 100%). To attain the desired degree of spatial aggregation, we built an R function that runs a Monte Carlo simulation. For a given number of land types, the simulation proposed a random change to the number of cluster germs for creating a neutral landscape mosaic using the Gibbs algorithm (Gaucherel, 2008). The proposed change was proportional to the distance between current and target relative mutual information, until the target was achieved within a given tolerance (herein 2% distance from the desired percent aggregation). Since this function created a random realisation of reference map 1, we imposed an additional constraint to control for the evenness of the land type cover. We had to control for the evenness of the land cover in map 1, because the following landscape parameter of land cover change controlled for the evenness in map 2, and hence for the degree of landscape change from simulated map 1 to map 2. For the measure of evenness, we used the normalised entropy, i.e. the Shannon entropy divided by its maximum value for the given number of land types (2% distance tolerance in the rescaled values with maximum of 100%). The normalised entropy, relative mutual information, and the neutral landscape mosaics were computed with the relevant functions of the respective R packages “asbio” (Aho, 2022), “landscapemetrics” (Hesselbarth et al., 2019), and “NLMR” (Sciaini et al., 2018). We focused on five different degrees of spatial aggregation in reference map 1: {10%, 30%, 50%, 70%, 90%}.

#### 3. The land cover change

This parameter controlled the evenness of land type cover in reference map 2. It was based on the normalised entropy of map 2. For constructing a demand distribution for map 2 with the desired normalised entropy, we built a function which ran another Monte Carlo simulation. In each iteration, the simulation proposed an allowed random increase in the cover of a randomly selected land type, and a respective decrease in another. If this proposal resulted to a value of normalised entropy closer to the target, it was accepted. The simulation ended when the desired normalised entropy was attained given some tolerance (2% distance from the target percent), with the additional constraint of all land types covering at least five pixels. Note that the parameter for the land cover change was 100% minus the percent normalised entropy. That is, one land type would cover the whole reference map 2 when the land cover change parameter was equal to 100%, whereas the distribution of land type cover in reference map 1 would be retained in map 2 when the land cover change parameter was 0%. The random realisations of simulated reference maps 2 would have five different degrees of normalised entropy in their demand distribution, i.e. with land cover change of: {10%, 30%, 50%, 70%, 90%}.

For comparing the performance of the models, we adopted a full factorial design with all combinations of the five values from the three parameters, leading to 5 × 5 × 5 = 125 combinations of land type count, spatial aggregation and land cover change. For each combination, we built *n* = 30 random realisations of reference landscapes, leading to 125 × 30 = 3750 pairs of reference maps 1–2. All simulated landscapes were on square grids of 100 × 100 cells.

Note that although the Monte Carlo simulation for the spatial aggregation parameter could create a reference map 1, the Monte Carlo simulation for the land cover change parameter could create only the demand for reference map 2. We hence needed to allocate this demand spatially, to complete the creation of a reference map 2. We used the “Ordered” allocation model (Fuchs et al., 2013). In that way, the model for creating the reference maps 2 was different from the compared models predicting map 2 in our tests of predictive performance. The Ordered model assumes a hierarchical order of socioeconomic value for the land types. We herein chose randomly this land type hierarchy in each landscape. The model focuses on each land type sequentially, in descending hierarchical order. If a focal land type’s demand in map 2 must be higher than that land type’s cover in map 1, the required cells from land types which are lower in the hierarchy, and with the highest suitability to the focal land type, are selected to become of the focal land type. If a focal land type’s demand is lower than in the previous time step, the required number of cells covered by the focal land type, and with the lowest suitability to the focal land type, become candidates for change to a hierarchically lower land type which will first require an increase in cover from map 1 to map 2. An R implementation of the Ordered model is provided in the R package “lulcc” (Moulds et al., 2015), and we herein used its non-stochastic, original version (Fuchs et al., 2013). Regarding the suitability, we simply employed logistic regression models relating the occurrence of each land type to a single environmental predictor. The map of the environmental predictor was a spatially correlated Gaussian random field with mean equal to 0.5 (0–1 value range), and high autocorrelation (autocorrelation range equal to the landscape’s side length, zero variation in the scale of the autocorrelation range, and magnitude of variation over the entire landscape equal to one), built with a relevant function from the R package “NLMR” (Sciaini et al., 2018).

We again here applied the CLUE-S and the trans-CLUE-S models to the simulated reference map 1 to predict map 2 under environmental or random suitability allocation, with none of the four model versions adopting any constraint in the five transition rules or in the elasticity settings. The models were based on the transition matrix of the reference maps 1–2. Additionally, we used the LUC of reference map 2 to build logistic models for suitability which are faster to build than random forest models given the large number of simulated landscapes, relating the occurrence of each land type to the single environmental predictor. Although it is recommended to avoid the use of reference map 2 for building the statistical models of suitability (Pontius et al., 2004), we did so mainly because we did not have data from previous time steps relating LUC with environmental conditions, because reference map 1 was built with the Monte Carlo simulation to accurately control the spatial aggregation and land type cover evenness. Only reference map 2 was built by using the single environmental predictor, i.e. with the environmental suitability component of the Ordered model. Since this same procedure was employed throughout the model comparisons on simulated reference landscapes, we expected the comparability of the results.

### 2.4. Quantifying the predictive performance of the models

Besides a visual assessment, we aimed for a more comprehensive, quantitative assessment of model predictive performance according to proposals in previous works (Pickard and Meentemeyer, 2019; Pontius et al., 2008). In specific, we analysed the reference maps 1–2 and the predicted map 2 of the empirical and simulated landscapes via their: (1) three-map comparison, for quantifying the allocation, quantity and total disagreement, together with the performance in respect to a null model; and (2) two-map comparison for the configuration disagreement.

The three-map comparison, contrary to a two-map comparison between only reference and predicted map 2, can distinguish the correctly predicted LUC change from the correctly predicted persistence, which is even more important in landscapes with lower LUC change (Pontius et al., 2011). Besides the two components of correct prediction (Fig. S1 in the Supplementary Material), that is, the correct change and correct persistence, the three-map comparison outputs the three components of error: (1) false change, i.e. reference persistence predicted as change; (2) wrong change, i.e. reference change predicted as change but to the wrong land type; and (3) false persistence, i.e. reference change predicted as persistence. These five components sum to the total cover of the study area.

About the related measures of disagreement (Fig. S1), we calculated the following three with the R package “lulcc” (Moulds et al., 2015):

#### 1. Allocation disagreement

This measure captures the proportion of cells with misallocated land cover (Chen and Pontius, 2010). If the predicted map 2 has land type quantities equal to the reference map 2, any disagreement between these maps would be due to misallocation of land cover (Fig. S1c). Half of the cells with misallocated land cover would be of false persistence, and the other half would be of false change. This disagreement could be completely resolved by spatially swapping the land types between the cells of false persistence versus false change. Allocation disagreement then is equal to the cover of cells which need to be swapped, i.e. two times the false persistence or false change (since the two components are equal). If one of the two error components is zero, there is no way to resolve the misallocation, i.e. allocation disagreement is zero (Fig. S1d). If the maps 2 have non-zero and unequal quantity in at least one of their land types, not all disagreement can be resolved via spatial swapping, i.e. false persistence and false change would be unequal (Fig. S1e). Allocation disagreement is then the overlapped error between the two components, multiplied by two for considering both halves of the cells with misallocated land types. In general, then, allocation disagreement is equal to two times the minimum between false change and false persistence: 2 min(*False change, False persistence*).

#### 2. Quantity disagreement

This measure captures the prediction’s disagreement in the quantity of land type transitions (Chen and Pontius, 2010). Quantity disagreement is the prediction error which would remain after spatially swapping the land types for resolving any misallocation (Fig. S1e). If one of the false persistence or false change is zero, there is an excess quantity of either false persistence or false change which is the quantity disagreement, even though allocation disagreement is zero (Fig. S1d). If both are non-zero but unequal, then there is an excess of irresolvable error from the larger error component. In general, then, quantity disagreement is the max(*False change, False persistence*) – min(*False change, False persistence*). Alternatively, it can be expressed as the absolute difference between false change and false persistence: |*False change* – *False persistence*|.

#### 3. Total disagreement

The total disagreement is the sum of allocation and quantity disagreement, plus the wrong change component of error (Varga et al., 2019).

Configuration disagreement quantifies the dissimilarity in the spatial patterning between reference and predicted map 2. This is important because the two maps can have the same amount of correctly predicted change, but may differ in the spatial arrangement of predicted change (Pickard and Meentemeyer, 2019). A common strategy for quantifying configuration disagreement is to first synthesise—for both the reference and the predicted map 2—their signatures of spatial patterning from a collection of landscape metrics, and consequently measure the distance between the signatures (Long et al., 2010; Pickard and Meentemeyer, 2019). However, signatures based on landscape metrics have been shown to suffer from issues such as the normalisation and the weighting of the individual landscape metrics, let alone the decision on which of the tens of available metrics to include, especially given their interrelations which are difficult to interpret (Niesterowicz and Stepinski, 2016). We hence used a more synthetic measure as a spatial signature, i.e. the co-occurrence distribution which is derived from a land type co-occurrence matrix (Nowosad and Stepinski, 2021). This discrete distribution captures the proportion of a map’s cells which co-occur adjacently for each paired combination of land types, including adjacent cells of the same land type. We built the two spatial signatures of the reference and predicted map 2, and quantified the dissimilarity between the two discrete distributions, with the 0–1 normalised measure of the robust Jensen-Shannon divergence, with the R package “motif” (Nowosad, 2021). A zero configuration disagreement means that the reference and predicted map 2 have identical co-occurrence distributions, whereas a unit disagreement means that they are so different that they do not share any common land types.

To compare model performance with a null model’s (Pontius et al., 2008), we can assume a naive model which predicts full persistence in map 2 from the reference map 1 (Fig. S2). In specific, such a null model scores a correct persistence equal to the observed persistence, and a false persistence equal to the observed change, while the rest of the components are equal to zero (Fig. S2c). Thus, a model which can predict change will perform better than null if it achieves more correctly predicted change than falsely predicted change (Fig. S2d,e).

For plotting purposes, we rescaled the 0–1 values of all measures of disagreement to the 0–100% interval. Whenever we had to present the disagreement from multiple replicate runs of a model under the same settings, we would either plot the mean disagreement (after confirming that the mean is representative by checking that the median is relatively close, and that the standard error is relatively small), or its distribution (with boxplots).

### 2.5. Comparing the sensitivity of models to the number of suitability predictors

For the empirical reference landscape, we ran a sensitivity analysis to compare the robustness of the CLUE-S and trans-CLUE-S predictions with environmental suitability allocation under different numbers of suitability predictors. Again here in this application on the empirical landscape, we furthermore tested the no-demand version for reference purposes, i.e. via simply predicting environmental suitability with the random forest model. We expected that the four measures of disagreement would score higher for fewer predictors. Was the allocation, and consequently the accuracy, of one model more influenced than the other by the amount of information about the environmental conditions of the study area?

Although the number of predictors in the empirical reference landscape was 13, we excluded the two categorical factors regarding the presence of two types of bedrock, to facilitate plausible land type classifications in map cells when the predictor number was low. For the remaining 11 predictors, we tried all numbers of predictors from one to 10, and in all combinations of predictors. That is, for one predictor, we tried each predictor separately, for a total of 11 runs; for two predictors, we tried all combinations of predictors in pairs, for a total of 55 runs; and for the rest of the predictor numbers, the total numbers of combinations were 165 (3 predictors), 330 (4), 462 (5), 462 (6), 330 (7), 165 (8), 55 (9), and 11 (10). Thus, the total number of model runs on the empirical landscape was 2046 for CLUE-S and 2046 for trans-CLUE-S. We applied the CLUE-S and trans-CLUE-S with all the combinations of suitability predictors, plotting each disagreement measure’s mean and standard error among the combinations of each number of predictors. The settings for the CLUE-S and trans-CLUE-S were the same as in their application to the original empirical reference landscape.

### 2.6. Comparing statistically the model predictive performance on simulated landscapes

We additionally aimed to statistically compare the predictive performance of the models with environmental suitability on the basis of the four measures of disagreement, and for the 3750 simulated landscapes built according to the three landscape parameters (number of land types, spatial aggregation and land cover change). For each measure of disagreement as a response variable, we built a statistical model relating the response disagreement to the three landscape parameters, together with the interaction of the model version with each landscape parameter: *disagreement* ∼ (*land type count + spatial aggregation + land cover change*) * *model*. Since the disagreement measures were bounded in the [0, 1] interval, we had to use beta regression models (Ferrari and Cribari-Neto, 2010). Beta regression accepts values in the open interval (0, 1), and we hence changed any 0 and 1 values to 0.00001 and 0.99999, respectively (Ferrari and Cribari-Neto, 2010). We formulated and ran the four beta regression models under maximum likelihood estimation with the R package “betareg” (Cribari-Neto and Zeileis, 2010). We assumed fixed dispersion across the values of landscape parameters, since variable dispersion did not improve model fit. Moreover, we used the “loglog” link function, instead of the default “logit”, because it is more appropriate under the presence of extreme values (Cribari-Neto and Zeileis, 2010). In our case, we had high frequency of disagreement values close to zero, and the “loglog” link function greatly improved model fit.

Since the estimated parameters from the beta regression models with “loglog” link function are difficult to interpret, we built effect plots to present the effect of the three landscape parameters on the four measures of disagreement. In specific, we plotted the 95% confidence limits of the mean effect of each landscape parameter under both model versions with the help of the R package “effects” (Fox et al., 2022).

## 3. Results

### 3.1. Comparing the models applied to the empirical reference landscape

A visual comparison of the predictions from four model versions revealed the higher similarity to the reference map 2 under higher resolution of demand, and under more informed suitability allocation (Fig. 2, from CLUE-S to trans-CLUE-S allocation, and from random to environmental suitability). For instance, the trans-CLUE-S model predicted the occurrence of specific patches, such as the grassland at the north (Fig. 2e), whereas the CLUE-S model predicted wrongly the occurrence of other patches, such as the open-scrub patch close to the centre (Fig. 2c).

**Fig. 2.**
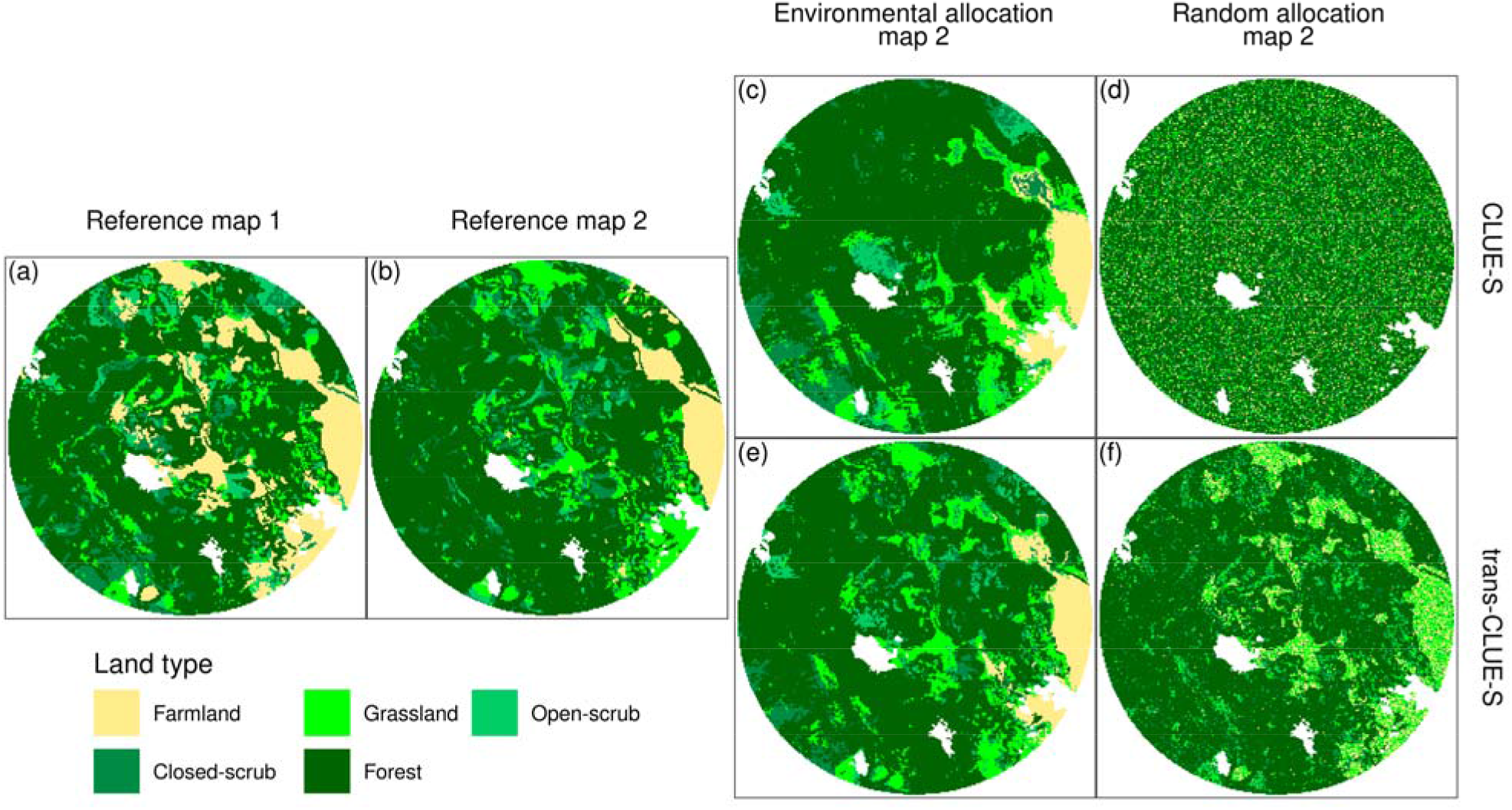
Model predictions of the empirical landscape. The empirical reference map 1 (a) and map 2 (b) are from years 1996 and 2015, respectively. The rest of the panels regard the prediction of reference map 2 by four model versions, with the type of demand resolution on the rows, and the type of suitability allocation on the columns, according to the side and top titles, respectively. Each random allocation map 2 is a single random realisation out of many possible (panels d and f). Map cells had 25 m side length (diameter of the circular study area is equal to 6 km, oriented vertically to the North).

Additionally, trans-CLUE-S retained some basic spatial features of reference map 2 under random allocation of demand (Fig. 2f), whereas CLUE-S returned an overall random spatial structure (Fig. 2d).

The measures of disagreement confirmed quantitatively this visual identification of higher performance under higher resolution of demand, and under more informed suitability allocation (Fig. 3). The reference version of no demand exhibited the worst performance under environmental suitability for all measures of disagreement, and under random suitability except of allocation disagreement. The CLUE-S model had slightly higher allocation disagreement than trans-CLUE-S, but much lower with random suitability, although their difference under environmental suitability was around 2% (Fig. 3a). Nevertheless, CLUE-S had also a similar amount of quantity disagreement as allocation disagreement under environmental suitability, and around double of that under random suitability. On the contrary, the trans-CLUE-S model had no quantity disagreement under both environmental and random suitability allocation (Fig. 3b). Adding the map percentage of wrong change to the previous two measures resulted in a total disagreement which was approximately one time higher for CLUE-S under both modes of suitability allocation (Fig. 3c). Finally, both models exhibited similar performance in the spatial characteristics of the predictions with environment-based allocation, but the random allocations of demand from CLUE-S had higher spatial disagreement with the reference map 2 than from trans-CLUE-S (Fig. 3d). Comparing the models to the null model naively assuming full persistence from map 1 to map 2, the trans-CLUE-S model performed better than null in all measures, whereas CLUE-S never performed better than null, under both environmental and random suitability allocation.

**Fig. 3.**
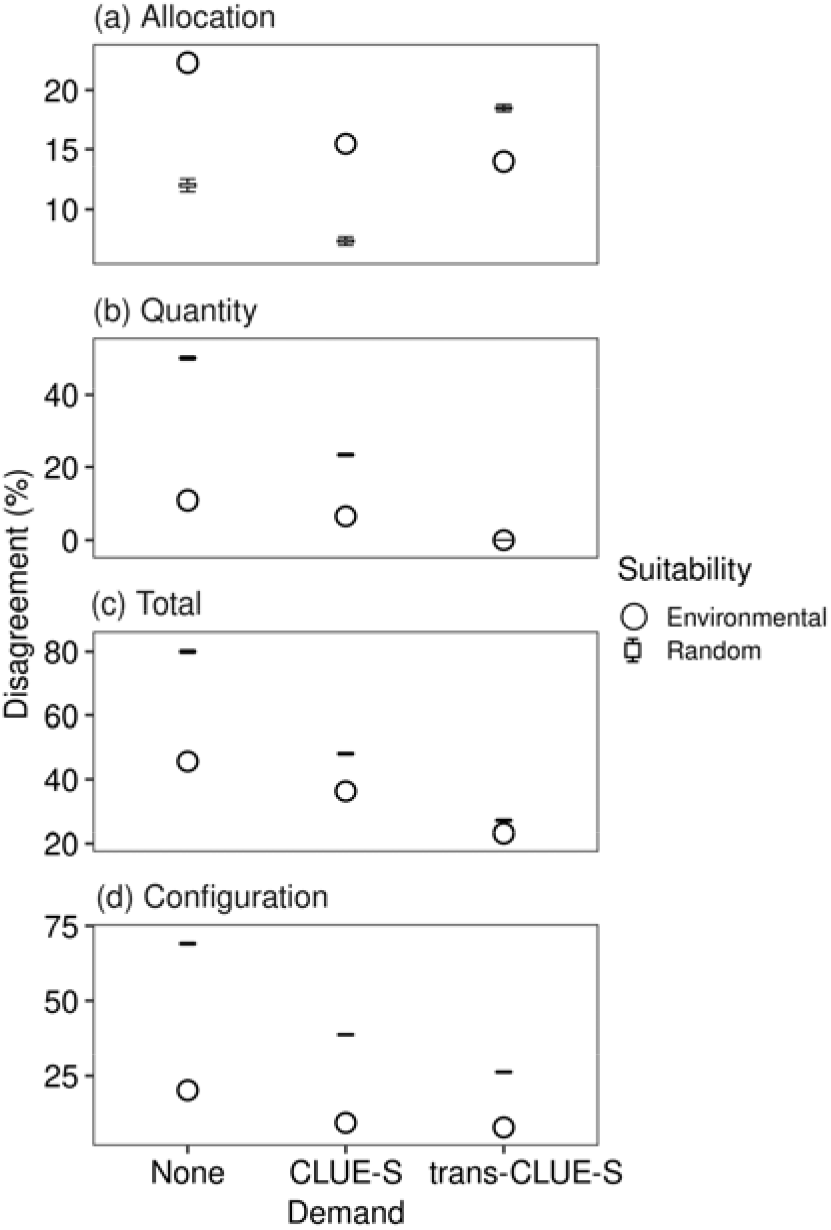
Predictive performance of six model versions applied to the empirical reference landscape. (a–c) These measures are based on the three-map comparison of the reference maps 1 and 2 and the predicted map 2. (d) Configuration disagreement is based on the spatial signature distance between reference and predicted map 2. For the three demand versions (x-axis), environmental suitability allocation is given with points. The same three demand versions but from 1000 random suitability allocations are given in boxplots (narrowly distributed, with outliers omitted because they were relatively close to the boxplots). The no demand version (None) produced mere suitability maps predicted by only the random forest model of suitability, i.e. without any demand restrictions to the number of cells covered by each land type.

We noted both visually (Fig. 2) and quantitatively (Fig. 3) that the performance of random allocations from trans-CLUE-S were closer to the allocations via environmental suitability than from the CLUE-S model. We hence elucidated further on these observations with simulations in the continuum between these two extremes of complete absence and presence of environmental predictors (Fig. 4). Indeed, the spatial similarity of the CLUE-S predictions to the reference maps 2 was more sensitive to this amount of environmental information, converging to its similar performance to trans-CLUE-S only when using most of the predictors (Fig. 4d). The highest spatial sensitivity to the number of environmental predictors was shown for the no-demand version of plain environmental suitability maps. For total disagreement, the trans-CLUE-S model exhibited more robust performance, since this measure increased more slowly under fewer environmental predictors than the CLUE-S and no-demand models (Fig. 4c). About the two components of total disagreement, the lower allocation disagreement of the CLUE-S and no-demand models decreased further for fewer environmental predictors, whereas it increased for trans-CLUE-S (Fig. 4a); and quantity disagreement increased for CLUE-S, and even more under no-demand, whereas it was zero throughout for trans-CLUE-S (Fig. 4b). Compared to the null (naive) model, the worst trans-CLUE-S performance was observed when 99.4% of the combinations performed better than null (it was in the case of three predictors), whereas the performance of CLUE-S was never higher than null under any number of predictors.

**Fig. 4.**
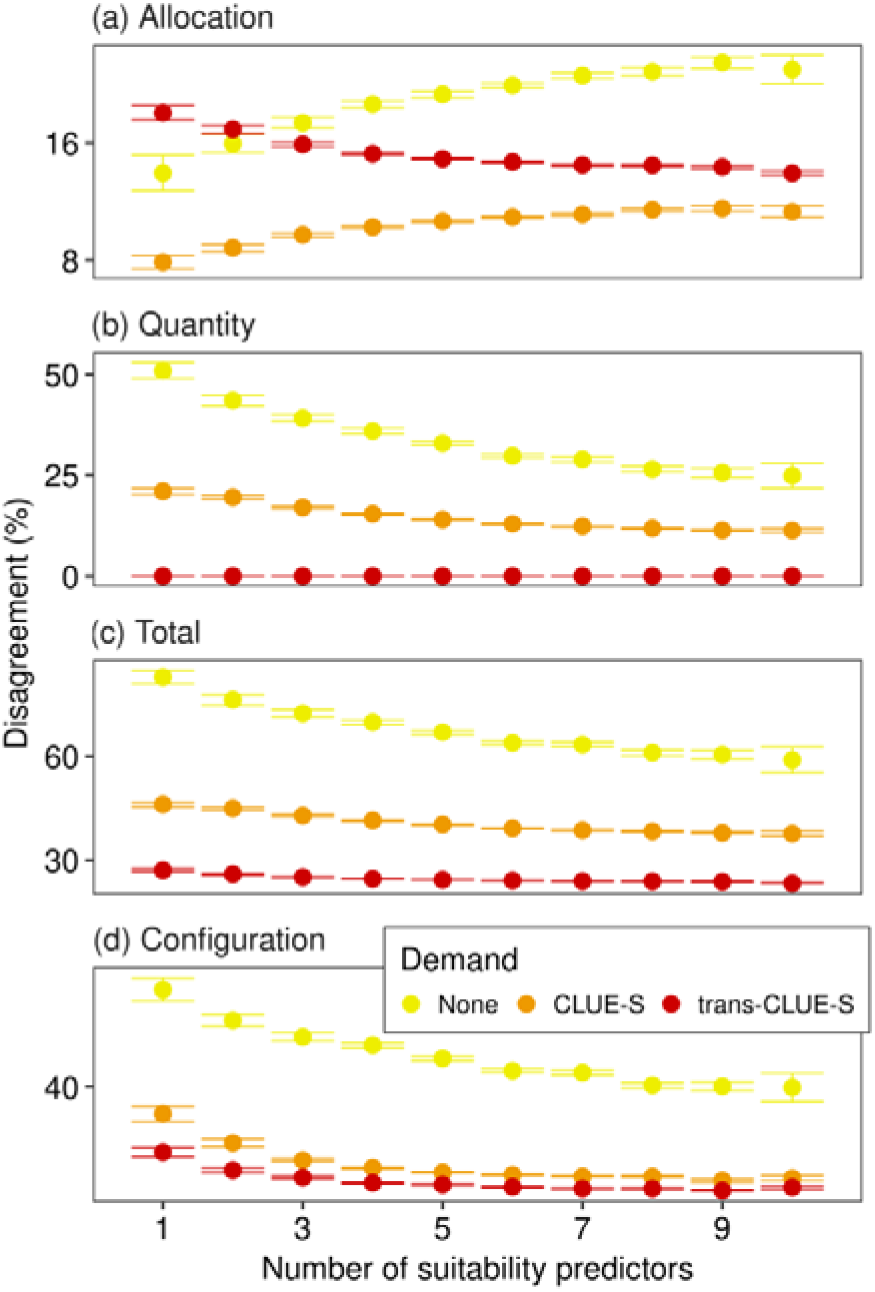
Sensitivity of CLUE-S and trans-CLUE-S performance to the number of environmental suitability predictors (empirical reference landscape). For each number of suitability predictors, all possible combinations from 11 predictors were tried in the random forest modelling of suitability. The points are the mean disagreement from all possible combinations of predictors (± the standard error). For reference, we additionally provide the model version of no demand which returned environmental suitability maps predicted only by the random forest model.

### 3.2. Comparing the models applied to simulated reference landscapes

A visual comparison of predictions from CLUE-S and trans-CLUE-S which were applied to simulated reference landscapes of contrasting characteristics reveals the more accurate reproduction of reference maps 2 by trans-CLUE-S (Fig. 5). Although the CLUE-S model managed to reproduce some basic features of reference map 2, it tended to allocate demand in a spatially more concentrated way, corresponding to the aggregated spatial distribution of the single environmental predictor. This was evident in both a highly aggregated and changing reference landscape of three land types (Fig. 5c), as well as in a highly disaggregated and stable in net change landscape of 15 land types (Fig. 5g). On the contrary, the trans-CLUE-S model managed to predict the position and shape of even small patches in both examples of reference landscapes (Fig. 5d,h).

**Fig. 5.**
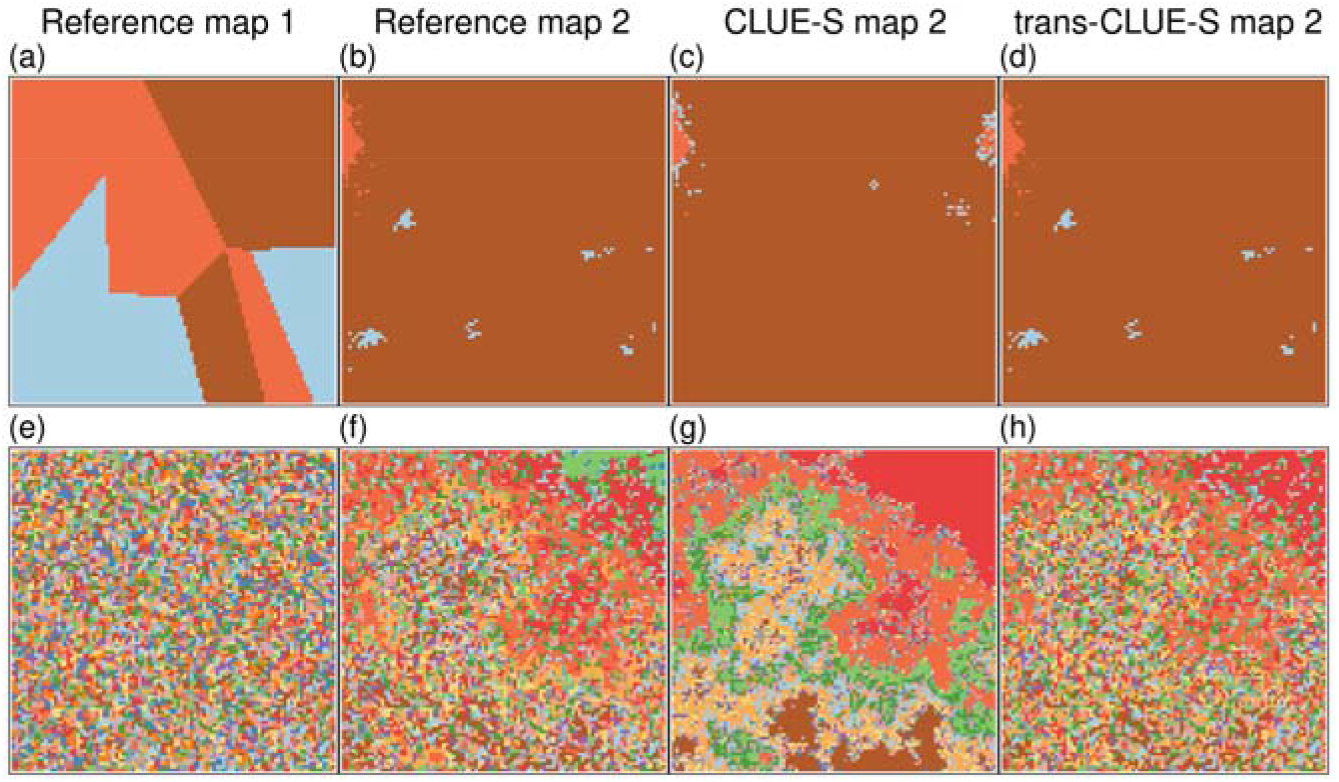
Indicative model predictions for two cases of simulated reference landscapes of contrasting characteristics. The first case in the top row of panels depicts: (a) a simulated reference map 1 with three land types and 90% spatial aggregation; (b) 90% land cover change in map 2; and (c,d) the predictions of map 2 from the CLUE-S and trans-CLUE-S model, respectively. The second case in the bottom row of panels depicts: (e) a simulated reference map 1 with 15 land types and 10% spatial aggregation; (f) 10% land cover change in map 2; and (g,h) the predictions of map 2 from the CLUE-S and trans-CLUE-S model, respectively. Land type cover was approximately even among the land types of maps 1. All maps had a size of 100 × 100 cells.

As indicated by the visual inspection, the predictive performance of the trans-CLUE-S model was commonly found quantitatively higher than CLUE-S when applied to simulated reference landscapes of a wide range of characteristics (e.g. see Fig. 6 for nine land types). For any number of land types, and in decreasing order of absolute difference in performance between the models, the median disagreement for respectively the trans-CLUE-S and CLUE-S were 7.8% and 28.2% for total disagreement, 0% and 15.6% for quantity disagreement, 10.4% and 18.2% for configuration disagreement, and 4.5% and 1.1% for allocation disagreement. Both models appeared to perform better in reference landscapes which were more spatially aggregated and which changed more between maps 1–2, although spatial aggregation seemed to have a weaker relation with model performance (e.g. see Fig. 6). Similar results were found for the other numbers of land types (Figs. S3–S6). Compared to the null model of full persistence, trans-CLUE-S performed better than null in 99.7% of the landscapes, whereas CLUE-S performed better than null in 77.6% of the landscapes, for any given combination of reference landscape parameters (number of land types included).

**Fig. 6.**
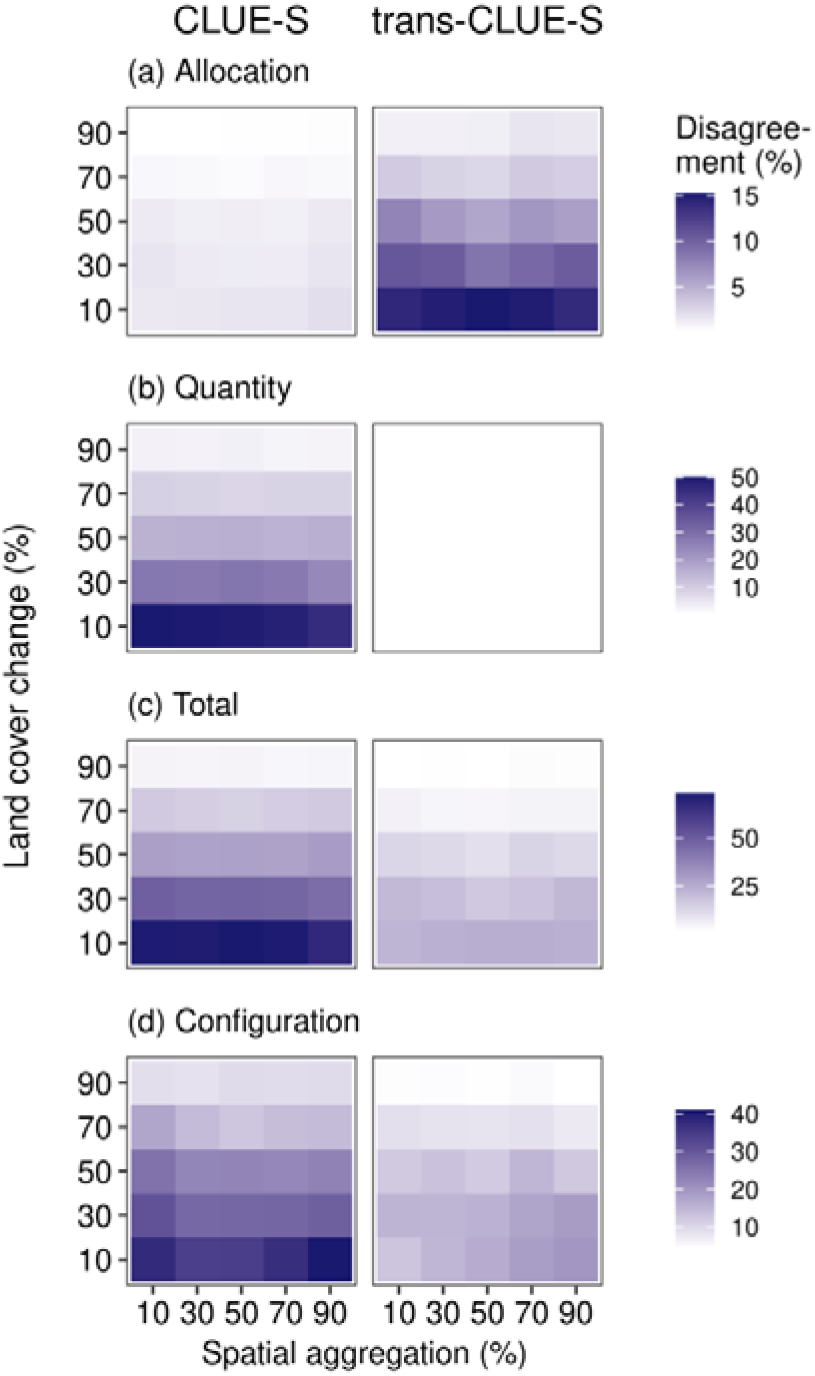
Predictive performance of CLUE-S (left column) and trans-CLUE-S (right) applied to simulated reference landscapes of different characteristics. Each pair of reference maps 1 and 2 was simulated on the basis of three parameters: number of land types in both maps (nine for this figure), spatial aggregation of map 1 (x-axis), and land cover change realised on map 2 (y-axis). For each combination of spatial aggregation and land cover change, CLUE-S and trans-CLUE-S were applied to the same simulated reference landscapes (*n* = 30 pairs of reference maps 1 and 2), here showing the mean percent disagreement.

The effects of all three parameters of the simulated reference landscapes could be more comprehensively investigated with the beta regression modelling (Fig. 7). These marginal effects similarly showed that although CLUE-S had lower allocation disagreement, it overall performed worse than trans-CLUE-S. In order of decreasing influence of the landscape parameters, it was confirmed that both models performed better under greater land cover change, fewer land types, and higher spatial aggregation. In other words, both models performed worst in landscapes covered by many land types, and for which the approximately even distribution of land type cover in map 1 changed minimally in map 2 (contrasting examples already shown in Fig. 5). Commonly, the model with worse overall performance (higher intercept) improved faster (steeper slope). Regarding the statistical significance, the effects of the landscape parameters, of the model type, and of the interaction between each parameter with model type were almost all significant according to the beta regression modelling (*p* < 0.001 for all four measures of disagreement). The only exception was the statistical insignificance of spatial aggregation on allocation disagreement (Fig. 7b). The four beta regression fits had a pseudo-R^2^ of 54%, 89%, 82% and 64% for allocation, quantity, total and configuration disagreement, respectively.

**Fig. 7.**
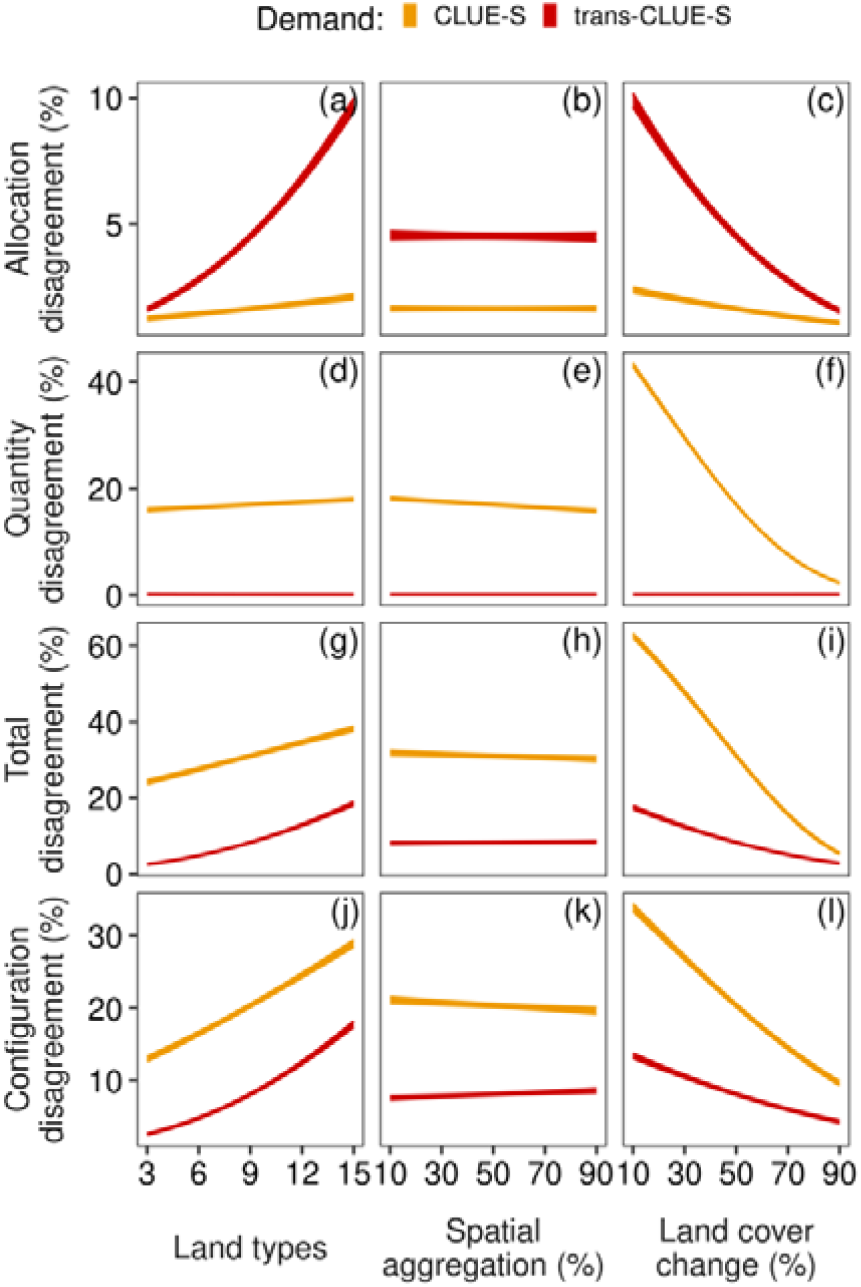
Marginal effects of the simulated reference landscape parameters on the predictive performance of CLUE-S and trans-CLUE-S. The shaded envelope regions are 95% confidence limits of the mean marginal effects based on a beta regression for each measure of disagreement. The beta regressions included the interaction between model type and each landscape parameter, hence the varying slope between CLUE-S and trans-CLUE-S.

## 4. Discussion

Although a popular choice for modelling LUC change, the CLUE-S model attempts to satisfy the demand for only the total cover of each land type in the predicted map 2. In the present work, we introduced new variants of CLUE-S with different resolutions of demand, one of which attempts to satisfy demand at the more detailed level of land type transitions, similarly to other spatial models. We found that the predictions of this trans-CLUE-S model had half the total and configuration disagreement of the CLUE-S predictions based on environmental suitability for both our empirical landscape and for numerous simulated landscapes of different characteristics. Regarding random or less informed suitability, trans-CLUE-S performed overall better than CLUE-S, hence exhibiting lower sensitivity to the amount of environmental information used for the spatial allocation of demand. In the following discussion, we attempt to explain the results from the model comparisons, and the advantages and disadvantages of the different models and of our methodologies.

Visual inspection of the predictions indicated that trans-CLUE-S was more able to reproduce fine features of the reference map 2 than CLUE-S. Despite its subjective nature, visual inspection of predicted maps can provide a preliminary and complementary assessment to quantitative assessment, identifying features which are not easily captured by objective measures of accuracy (García-Álvarez et al., 2019). For instance, we noted that trans-CLUE-S successfully predicted the occurrence of the grassland patch at the north of the empirical landscape, whereas CLUE-S missed it (Fig. 2). This happened because trans-CLUE-S focuses on the cells of each land type in map 1 separately. The northern patch was farmland in map 1 before turning to grassland in map 2. Since trans-CLUE-S focused on the farmland cells of map 1, it was more likely to turn with higher spatial accuracy that patch to grassland via the environmental suitability allocation. The CLUE-S model did not have the farmland pixels of map 1 as basis, and thus allocated the demanded grassland anywhere suitable in the whole landscape. For the same reason, trans-CLUE-S was also more successful under random or less informed suitability. In specific, allocation of demand was random or less informed only inside the cells of each land type, whereas CLUE-S applied such an allocation to the whole landscape, resulting in a spatially random distribution of demand. We could reduce this kind of CLUE-S error by employing transition rules and elasticity settings, but this would require extensive manual parameterisation via trial and error (Mas et al., 2014). On contrary, our trans-CLUE-S extension is algorithmic, i.e. without introducing new parameters, hence reducing the need to employ the two features of transition rules and elasticity.

This employment of map 1 by trans-CLUE-S is characteristic of other dynamic, spatial, inductive and pattern-based models. An example is the Ordered model which we herein used for the creation of simulated reference landscapes. Despite the adoption of demand at the level of total land type cover for map 2, similarly to the CLUE-S model, the Ordered model starts with map 1 and alters it according to suitability until demand is met (Fuchs et al., 2013). The reason that the Ordered model can incorporate map 1 is the assumed hierarchy of land types, enabling the allocation routine to focus on the map 1 cells of each land type separately, instead of the whole landscape as in CLUE-S. As Fuchs et al. (2013) discuss about their model, the assumption of a land type hierarchy might not be always and everywhere plausible. On contrary, our trans-CLUE-S model can focus on the map 1 cells of each land type separately and in any order, without the need of a hierarchy assumption. This is possible because trans-CLUE-S works with the transitions of each land type. Other popular models of the same category are similarly based on map 1, but mainly because they are cellular automata, such as the CA-Markov and DINAMICA models (Paegelow and Olmedo, 2005; Soares-Filho et al., 2002). Such cellular automata will be based on map 1 to re-weight the probability of LUC change according to near-neighbourhood or far-off rules, which is also possible with the original CLUE-S model by adopting the neighbourhood transition rules. The differentiating feature of our trans-CLUE-S model, though, is that it exploits all the information of the transition matrix both non-spatially (quantity) and spatially (allocation), i.e. it processes directly from map 1 the proportion of a land type’s cells which will persist or change to any other land type.

Non-spatially, the trans-CLUE-S usage of all the information from the transition matrix was in part responsible for its higher predictive performance. The trans-CLUE-S model exhibited zero quantity disagreement because we used the realised transitions from the transition matrix of the reference maps 1–2. In that way, the same land cover which was predicted falsely as changing was somewhere else in the landscape predicted falsely as persistent. As Pontius et al. (2004) recommend, reference map 2 should not be used during the parameterisation phase. In another work though, Pontius et al. (2008) show that accuracy assessments can be confounded by quantity disagreement. Since the present work’s aim was the comparison of models, and not the development of an empirical model for a real-world application, we decided to minimise quantity disagreement by parameterising according to the reference map 2. The CLUE-S model had non-zero quantity disagreement because false persistence was not equal to false change, since only land type totals were taken into account, and not the realised transitions. As expected, the worst performance was exhibited by the no-demand model, showcasing the importance of employing the CLUE-S and trans-CLUE-S allocation routines for better LUC predictions than the ones produced by mere suitability maps.

Spatially, the aforementioned focus of trans-CLUE-S on the cells of each land type in map 1 was also responsible for higher predictive performance. This was evident from the inequality between false change and false persistence in CLUE-S: false change was much larger than false persistence, with the latter being similar to the false persistence of trans-CLUE-S. Although this resulted in similar allocation disagreement (similar misallocation could be resolved via swapping), CLUE-S predicted more spatially shifting landscapes than in real, given that it met completely the demand of total cover of all land types. This was even more pronounced under less informed suitability, i.e. with fewer environmental suitability predictors, where false persistence from CLUE-S decreased while false change increased, as allocation in the predicted maps 2 became more spatially diffused and random.

This resulted to further decrease in allocation disagreement (pairing of false change–persistence to resolve misallocation), and to further increase in quantity disagreement. Thus, this analysis provides another reason for the advantage of the more spatially concentrated allocation of the trans-CLUE-S model. Even under scarcer environmental information for allocation, focusing on the cells of each land type from map 1 reduced false change, which together with wrong change were the main constraints to the performance of CLUE-S.

Despite these differences between CLUE-S and trans-CLUE-S, both models performed worse in simulated landscapes of smaller land cover change and more land types. The positive relation of performance with land cover change has been identified in other applications of different models to empirical landscapes (Pontius et al., 2008). One reason that these authors suggest, and which seems plausible in our work, is that this relation arises because smaller net change might be accompanied by large enough swap change, which is more difficult to predict. This was evident in our simulated reference landscapes that had smaller net but larger swap change for smaller values of the land cover change parameter. Regarding the number of land types, Pontius et al. (2008) did not find any other strong effect on performance in their empirical landscapes. Nevertheless, our systematic and extensive testing on numerous simulated landscapes of different characteristics identified a statistically significant negative effect of the number of land types on performance. Similarly, Varga et al. (2020) have found that a decrease in the number of land types because of land type aggregation was related to higher predictive accuracy of the CA-Markov model. For a fixed degree of land cover change, we expect that more land types create more opportunities for increasing the error in different components of a model, such as in the non-spatial demand for each land type or land type transition, in the statistical model for the cell suitability to each land type, and the spatial allocation of demand for each land type. Finally, regarding the spatial aggregation of reference map 1, we believe that it did not have a strong relation with accuracy for different reasons between the two models. On one hand, the CLUE-S model did not take into account map 1, hence spatial aggregation did not matter. On the other hand, by focusing on the cells and hence on the patches of each land type separately, the trans-CLUE-S performance was not considerably affected by the patchiness of map 1.

One of the drawbacks of the trans-CLUE-S model is related to this focus on the map 1 cells of each land type, despite the provided advantage in predictive performance. Since trans-CLUE-S essentially executes a separate CLUE-S allocation for each land type, it requires more execution time to meet demand, or at least to converge to a desired distance from it. Moreover, as convergence issues can arise in CLUE-S, these issues can be on average multiplied by the number of land types in trans-CLUE-S. Such a frequent issue was the inability to satisfy demand at a desired distance due to cyclic non-convergence in our application of the models to the thousands of empirical landscapes with different numbers of environmental predictors, and of the simulated landscapes as well.

Alternatively, even if convergence was feasible, it would frequently require a prohibitive amount of time to reach small enough distance. Reaching small distance to demand, ideally satisfying demand completely, was important because the amount of predicted change could confound our accuracy assessment during the comparisons of model predictive performance, besides the obvious reason of increasing the accuracy of any empirical model for a real-world application (Pontius et al., 2018). These convergence challenges, and the aim of optimising the execution of CLUE-S and trans-CLUE-S, have led us to adopt and append the college admissions problem after reaching some distance from demand under the CLUE-S allocation routine. The college admissions problem could be solved relatively fast with the Gale–Sapley algorithm, which increases its execution time linearly with the size of the input data (Gale and Shapley, 1962). Taking over the CLUE-S loop after reaching a distance from demand of a few hundreds or thousands of cells for any land type, the college admissions execution would complete the full meeting of demand in seconds in our home computers (Tilly and Janetos, 2021).

Based on CLUE-S, the present study offered models with four variants of demand resolution, and two variants of suitability, leading to eight model versions (herein comparing only the six of them). The reason we developed all these model versions was for better distinguishing the contribution of individual model components. Regarding demand, the performance contribution of demand resolution can be distinguished by comparing predictions from different demand resolutions. For example, the performance contribution of demand at the level of land type transitions can be revealed by comparing the accuracy of the predicted map with the accuracy of a map predicted with no demand, as we did for reference in the comparisons of the empirical landscape, and in a first application of the trans-CLUE-S model for predicting empirical LUC in relation to climate change scenarios for year 2055 (Kiziridis et al., 2023). Regarding suitability, the comparison between maps predicted with environmental versus random suitability allocation can help us untangle the contributions to model predictive performance of the environmental information versus the allocation algorithm *per se*. As an example from the present study, such a comparison between environmental versus random allocation has led us to develop further simulations which identified CLUE-S as the model which was more vulnerable to the scarcity of environmental information for suitability-based allocation.

Due to limited space, we had to leave out the detailed presentation and comparison of all eight model versions. Preliminary comparisons between all model versions showed that the trans-CLUE-S model with its most detailed level of demand resolution outperformed the other three demand versions. This is the reason we herein focused on the comparison between this best of our new variants, versus its parental model CLUE-S. An opportunity for future investigations would be the comparison of all models, and in simulated reference landscapes with additional landscape parameters. For instance, we herein used simulated reference maps 1 with approximately even distribution of land type cover, but future comparisons can use different degrees of evenness. In that way, model behaviour can be further elucidated in important dimensions of landscape characteristics. Finally, an interesting future exercise would focus on the comparison between maps predicted solely via the CLUE-S allocation routine versus maps produced after appending the college admissions problem. Preliminary tests during the development of this new method showed high similarity with the results of the pure CLUE-S allocation. We believe that the great facilitation and acceleration of demand convergence provided by the addition of the college admissions allocation routine deserves a more systematic investigation.

## 5. Conclusions

The present work’s introduced CLUE-S variants lead to three main implications regarding the popular CLUE-S model for which the demand at the level of land type sums could be easily upgraded to the level of land type transitions. The first implication is the enhancement of the predictive ability of our toolkit with the addition of the trans-CLUE-S model which at the same time does not require considerably more computational resources than its parental CLUE-S model. This is because trans-CLUE-S showed higher predictive performance than CLUE-S, being able to predict even specific patches in the LUC map, which is especially relevant for the small scales we are commonly challenged to work with. Second, our predictions with trans-CLUE-S model are more robust under the commonly encountered scarcity of biophysical and especially socioeconomic data for suitability. This is a significant issue for future projections even for biophysical data, e.g. climatic scenarios for the future, because they commonly show high inter-model variability, but we showed that trans-CLUE-S predictions were less sensitive to the amount of such environmental information. Third, our model variants are not black-box models, but rather open-source projects which encourage the usability, extensibility, shareability and reproducibility of LUC research. The potential of the field to inform the public opinion, and the development of LUC policies, given specific climatic and socioeconomic scenarios for the future, can be further increased by shifting to a more transparent paradigm of LUC change research.

## Data accessibility

The R code for the models and auxiliary functions are uploaded to figshare (https://doi.org/10.6084/m9.figshare.21014869.v1).

## Supplementary material

All Supplementary Material can be found online for this article.

## Author contributions

Conceptualization, D.A.K. and I.T.; data curation, D.A.K., A.M., M.P., F.X. and I.T.; formal analysis, D.A.K.; investigation, D.A.K. and I.T.; methodology, D.A.K. and I.T.; software, D.A.K.; visualization, D.A.K.; writing—original draft, D.A.K.; writing—review and editing, D.A.K., A.M., M.P., S.T., F.X. and I.T.; supervision, F.X. and I.T.; funding acquisition, I.T.; project administration, I.T. All authors have read and agreed to the published version of the manuscript.

## Acknowledgements

The present study was supported by the Hellenic Foundation for Research and Innovation (H.F.R.I.) under the “1st Call for H.F.R.I. Research Projects to support Faculty Members & Researchers and the Procurement of High-cost Research Equipment Grant” [Project Number: 2333]. We acknowledge the Hellenic Cadastre, the Hellenic Military Geographical Service, and the Ministry of Rural Development and Food of the Hellenic Republic for providing orthophotos and large-scale aerial photographs. We would like to thank Georgia Bourdanou for the entry of socio-economic data, Grigorios Vassilopoulos for organising land cover mapping, Anastasios Zotos for gathering socio-economic data, and Ioannis Kokkoris for implementing orthorectification. We finally thank two anonymous reviewers for their insightful feedback which significantly improved the present work.

