## Supplementary Material (Figures S1--S6) for "Improving the predictive performance of CLUE-S by extending demand to land transitions: the trans-CLUE-S model"

by

Diogenis A. Kiziridis, Anna Mastrogianni, Magdalini Pleniou, Spyros Tsiftsis, Fotios Xystrakis, and Ioannis Tsiripidis

#### Figure S1: Three examples of disagreement metrics

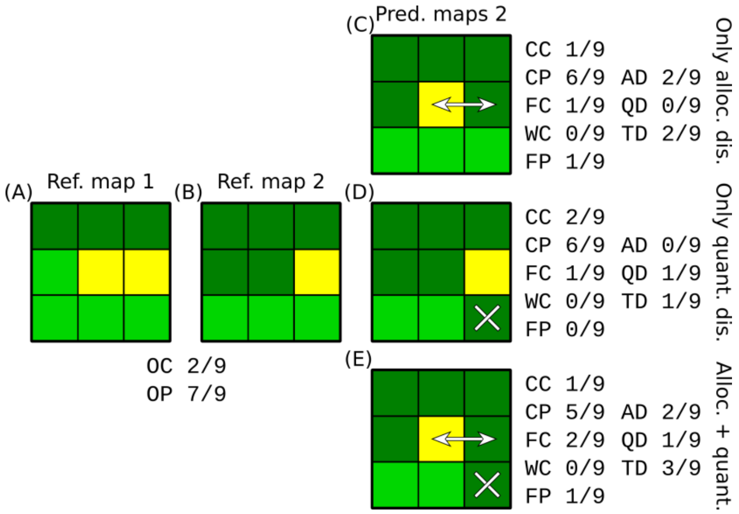

Fig. S1. Three examples of disagreement between reference and predicted map 2. The 3 × 3 landscape could have up to three land types (different colour and shading). The observed change (OC) from reference map 1 (A) to map 2 (B) was 2/9 of the landscape, and the observed persistence (OP) was the remaining 7/9. The five components of the three-map comparisons are the abbreviated: correct change (CC); correct persistence (CP); false change (FC); wrong change (WC); and false persistence (FP). The three examples of disagreement are: (C) only allocation disagreement (AD), because all disagreement can be resolved by swapping the land cover of two pixels (double-headed arrow); (D) only quantity disagreement (QD), because disagreement cannot be resolved by any swapping (excess quantity of the darkest land type indicated by “x”); and (E) allocation and quantity disagreement. In all examples, total disagreement (TD) is the sum of allocation and quantity disagreement, plus the wrong change.

#### Figure S2: Three examples for the null model on a 3 × 3 landscape

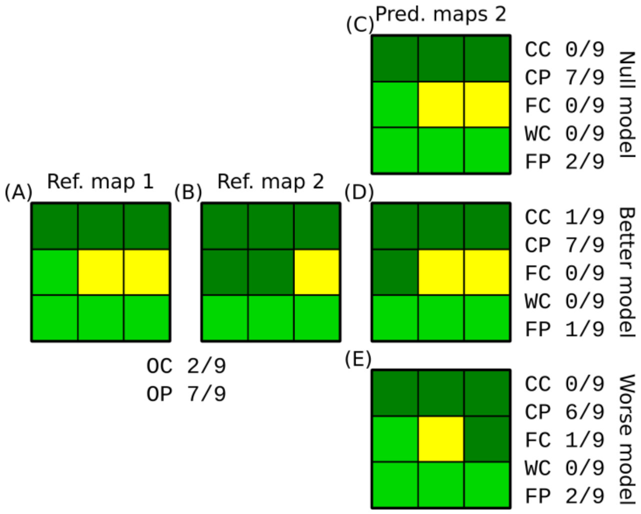

Fig. S2. Three examples for the null model on a 3 × 3 landscape with three land types of different colour and shading. The observed change (OC) from reference map 1 (A) to map 2 (B) was 2/9 of the landscape, and the observed persistence (OP) was the remaining 7/9. The five components of the three-map comparison are the abbreviated: correct change (CC); correct persistence (CP); false change (FC); wrong change (WC); and false persistence (FP). The three examples are: (C) the null model naively predicts that map 2 is the result of full persistence in reference map 1; (D) a model performing better than null predicts change correctly; and (E) a model performing worse than null predicts change where there was persistence.

#### Figure S3: Predictive performance of CLUE-S and trans-CLUE-S (3 land types)

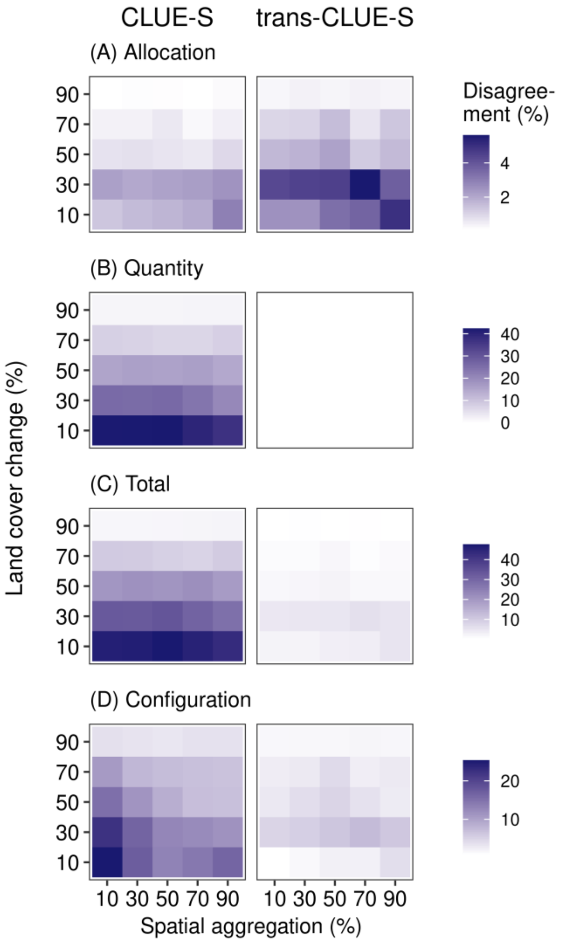

Fig. S3. Predictive performance of CLUE-S (left column) and trans-CLUE-S (right) applied to simulated reference landscapes of different characteristics. Each pair of reference maps 1 and 2 was simulated on the basis of three parameters: number of land types in both maps (three for this Figure), spatial aggregation of map 1 (x-axis), and land cover change realised on map 2 (y-axis). For each combination of spatial aggregation and land cover change, CLUE-S and trans-CLUE-S were applied to the same simulated reference landscapes ($n$ = 30 pairs of reference maps 1 and 2), here showing the mean percent disagreement.

### Figure S4: Predictive performance of CLUE-S and trans-CLUE-S (6 land types)

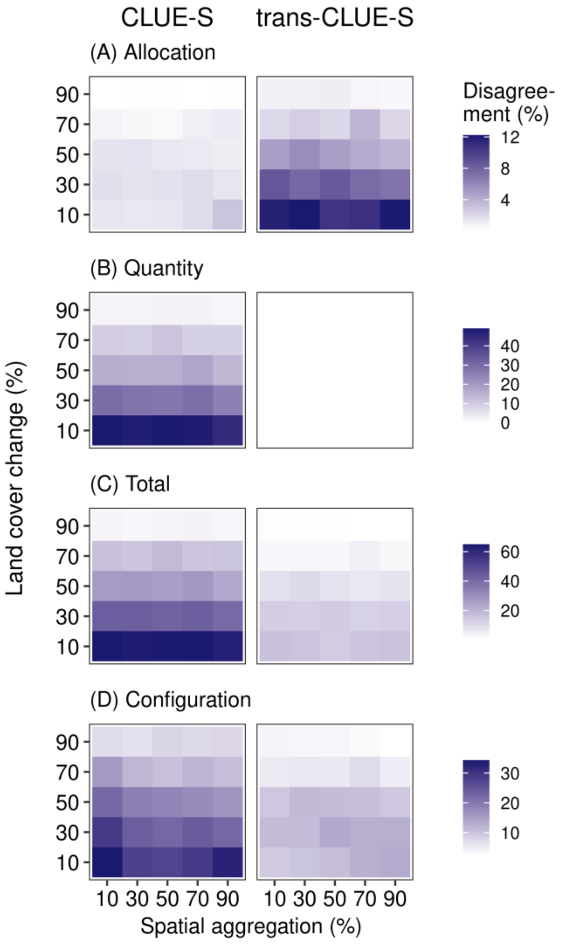

Fig. S4. Same as Fig. S3, but for six land types.

#### Figure S5: Predictive performance of CLUE-S and trans-CLUE-S (12 land types)

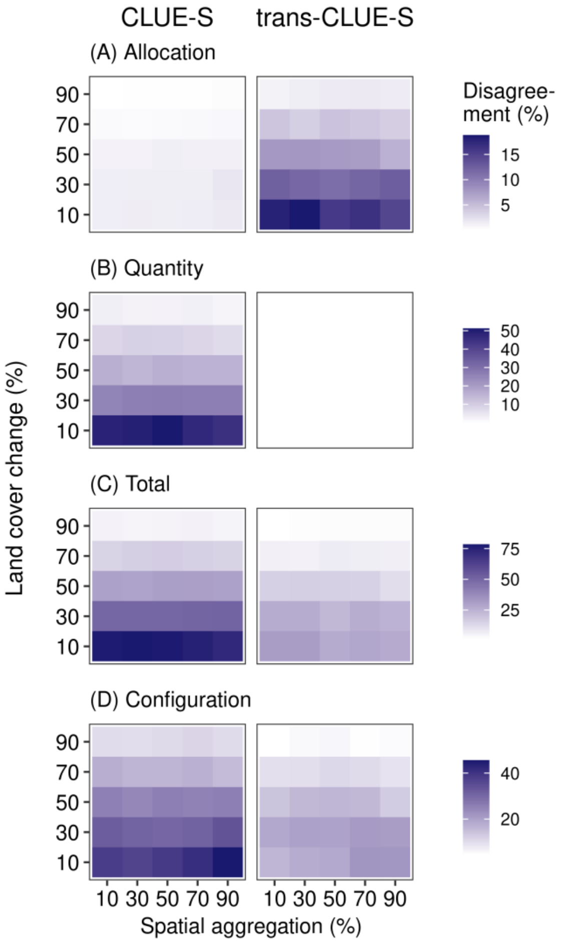

Fig. S5. Same as Fig. S3, but for 12 land types.

### Figure S6: Predictive performance of CLUE-S and trans-CLUE-S (15 land types)

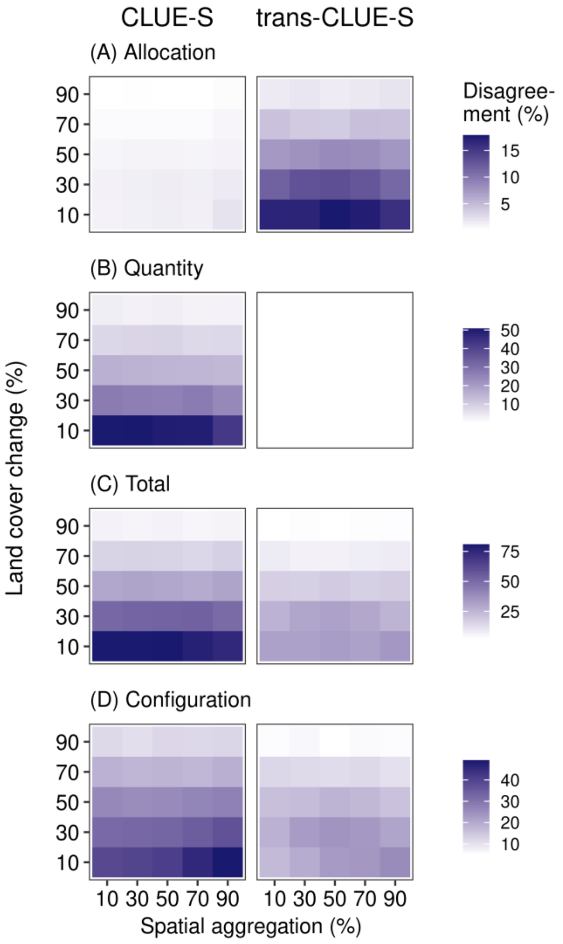

Fig. S6. Same as Fig. S3, but for 15 land types.
